## Supplementary material for "Developmental changes in story-evoked responses in the neocortex and hippocampus": figure supplement

### Supplementary Information

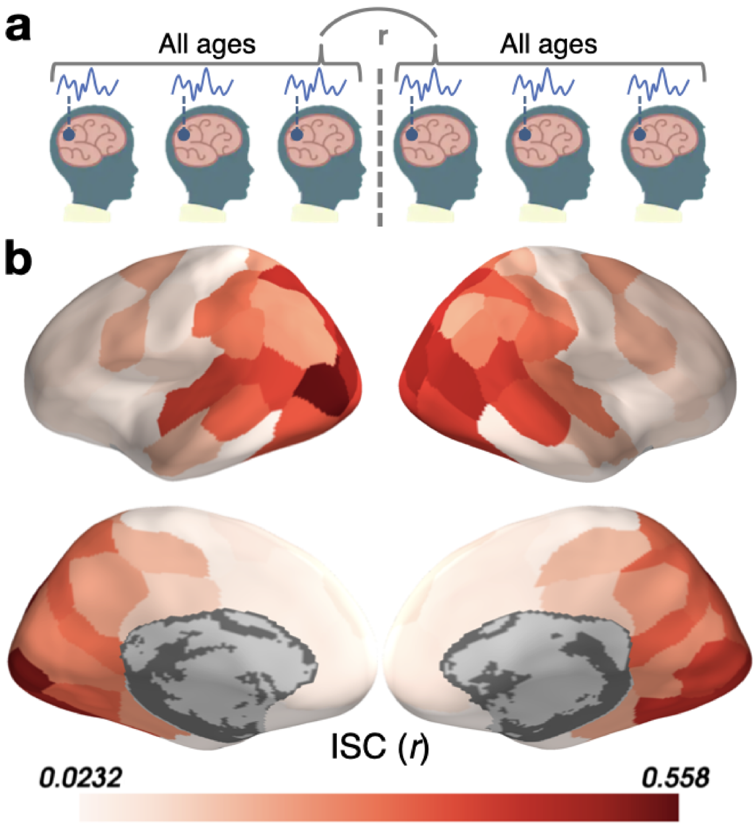


**Figure 1-figure supplement 1: Inter-subject correlation (ISC) across a random mixture of the Youngest and Oldest subjects.** (a) Schematic illustration of the correlation for each parcel between subjects in the Youngest (5-8 years) and Oldest (16-19 years) groups. (b) ISC in parcels on the cortical surface.

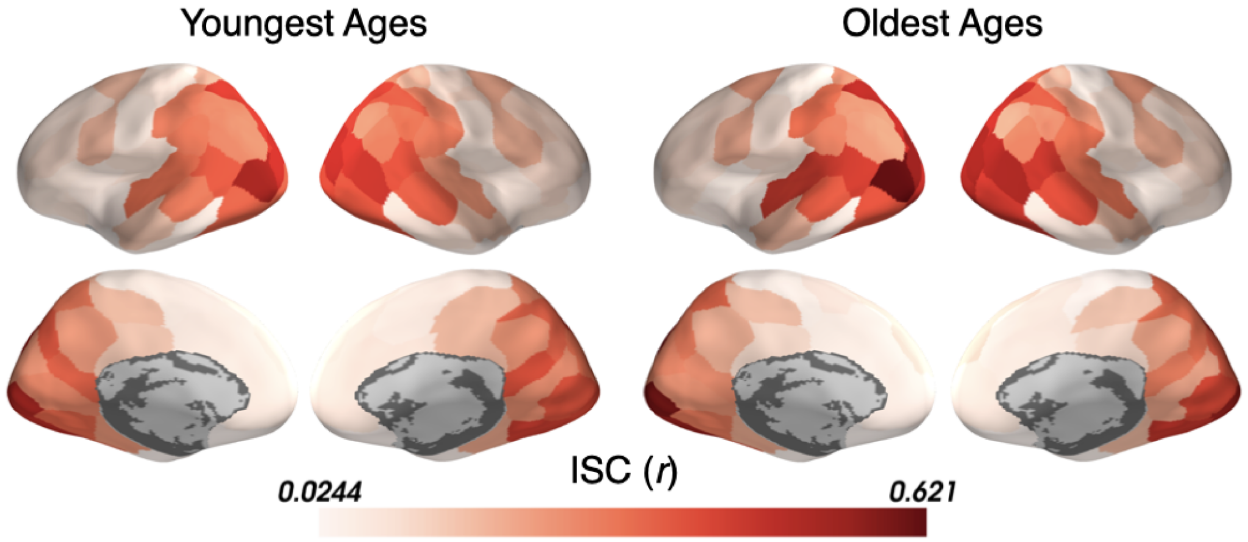


**Figure 1-figure supplement 2: Inter-subject correlation (ISC) within the Youngest and Oldest subjects.** ISC is displayed separately in parcels across the cortical surface for subjects in the Youngest (5-8 years) and Oldest (16-19 years) groups. Both age groups show a similar pattern of ISC increasing from sensory to anterior regions.

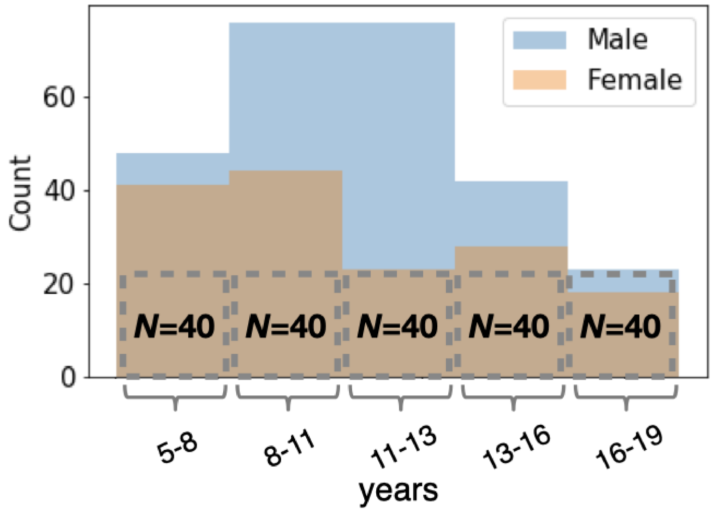


**Figure 1-figure supplement 3: The demographic breakdown of the subject groups studied.** The "Oldest" age group (16 - 19 years) had only 40 subjects. All other age groups contained a sub-sample of 40 subjects whose gender distribution matched that of the Oldest group. The counts of male and female subjects are overlapping.

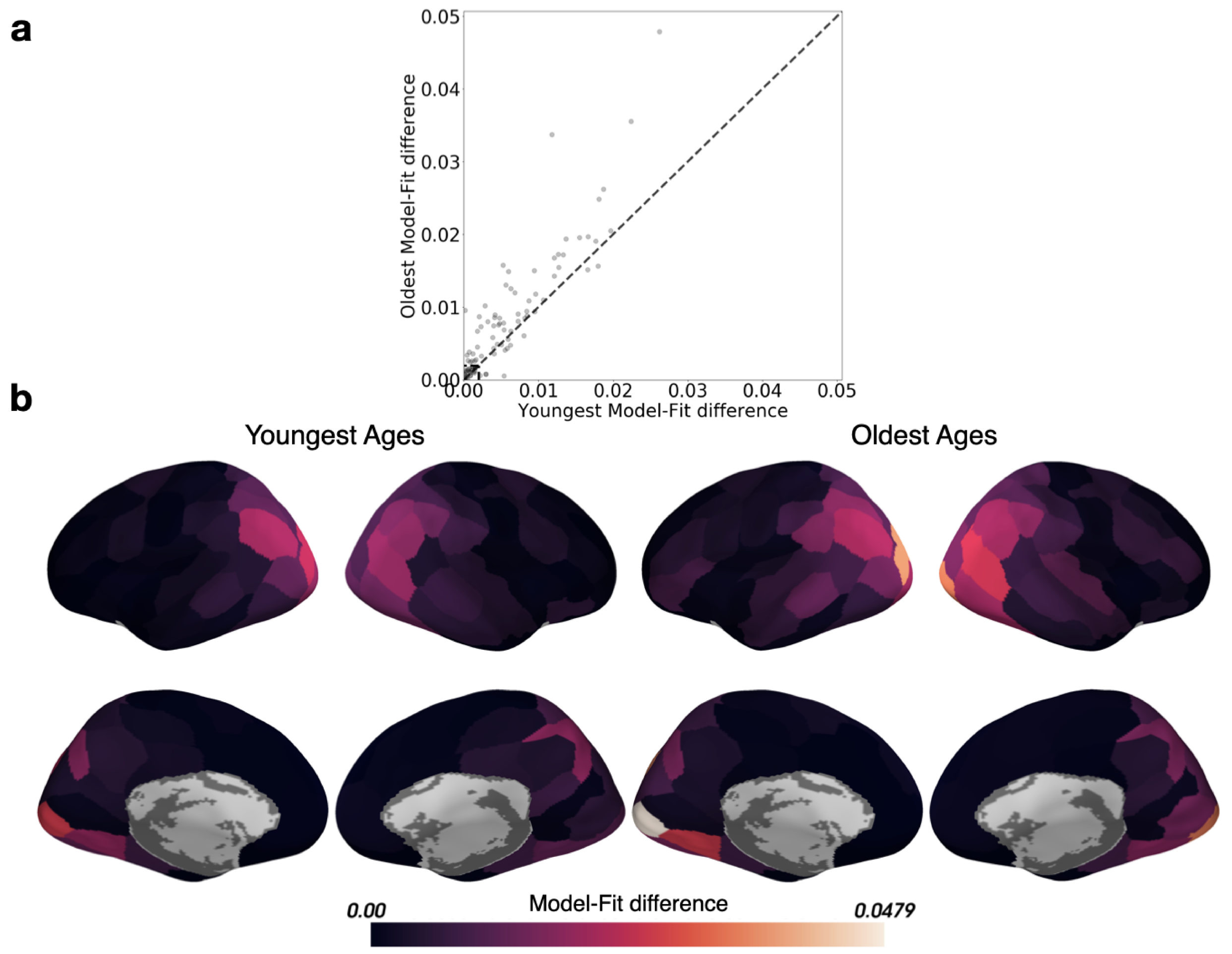


**Figure 5-figure supplement 1: Comparing the model-fit difference for all parcels between the Youngest and Oldest ages.** The model-fit difference is the difference in log-likelihood between the HMM for two events (expected to fit poorly), and the HMM with maximal model-fit (the best fitting number of events). (a) Parcels within the box in the lower left-hand corner of the scatter plot did not capture event structure in either age group (model fit differences smaller than 0.002), and were therefore excluded from the HMM-based analyses. (b) The model-fit difference is displayed in cortical parcels for both the Youngest and Oldest ages.

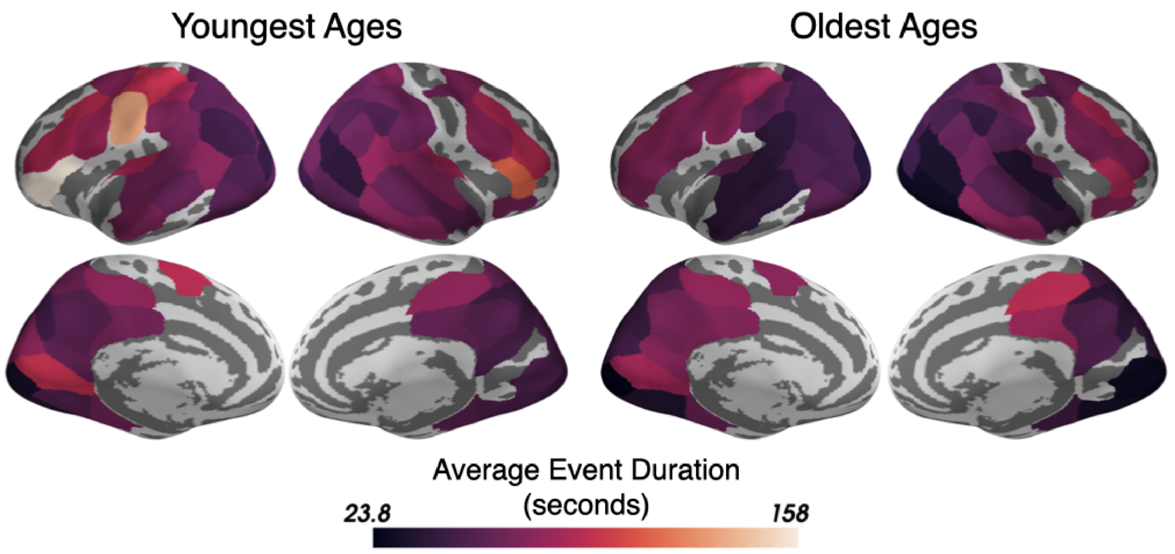


**Figure 5-figure supplement 2: The best-fitting average duration of events for the optimal HMMs trained and tested on either the Youngest or the Oldest ages.** The average event duration increases from sensory to association parcels for both age groups and the average event durations are highly correlated between the groups (r=0.78, RMS difference between groups = 12.3 seconds). Only parcels that had good model-fits in at least one age group (see Supplementary Figure 2) are shown.

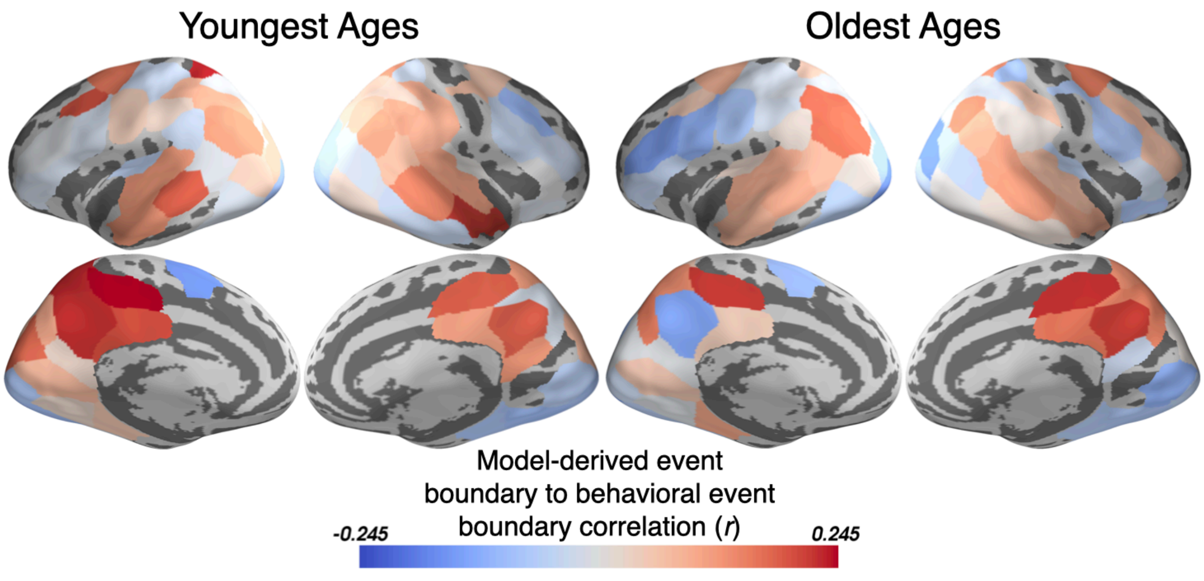


**Figure 5-figure supplement 3: The HMM-derived event boundaries correlate with behaviorally estimated event boundaries from children.** The event boundaries determined by the HMM jointly-fit to both the Youngest and Oldest groups correspond to behaviorally estimated event boundaries in association regions such as PMC, TPJ, and precuneus. All parcels for which a jointly fit HMM was modeled are displayed.
