## Appendix for "Developmental changes in story-evoked responses in the neocortex and hippocampus"

### Appendix A

For across-group analyses, we want to calculate the ISC between two groups of subjects, with two different sets of shared responses, $g_1$ and $g_2$, and different levels of variance, $\epsilon_i$ and $\epsilon_j$. To simplify the notation, we define $g_1^T g_1 = a_1$, $g_2^T g_2 = a_2$, $g_1^T g_2 = a_B$, $\mathbb{E}[\epsilon_i^T \epsilon_i] = b_1$ and $\mathbb{E}[\epsilon_j^T \epsilon_j] = b_2$. We would like to estimate the correlation between $g_1$ and $g_2$, regardless of the difference in signal-to-noise levels between the two groups. That is, our goal is to compute:

$$ISC_b = \frac{a_B}{\sqrt{a_1}\sqrt{a_2}}$$

If we know the ISCs for each group, from equation 1:

$$shISC_1 = \frac{N^2 a_1}{N^2 a_1 + 4N b_1}$$

and

$$shISC_2 = \frac{N^2 a_2}{N^2 a_2 + 4N b_2},$$

where $N$ is the number of subjects in each group, and the ISC between the two groups is:

$$shISC_B = \frac{N^2 a_B}{\sqrt{N^2 a_1 + 4N b_1}\sqrt{N^2 a_2 + 4N b_2}}.$$

Dividing $shISC_B$ by the geometric mean of $shISC_1$ and $shISC_2$ allows us to calculate the between-group ISC without the $b$'s:

$$\begin{aligned} ISC_b &= \frac{shISC_B}{\sqrt{shISC_1}\sqrt{shISC_2}} \\ &= \frac{a_B}{\sqrt{a_1}\sqrt{a_2}}. \end{aligned}$$

### Appendix B

We start by assuming that the response timecourse for every subject consists of a response that is shared across subjects, $g$, and a subject-specific response, $\epsilon$. The response timecourse for subject $i$ can therefore be represented as $s_i = g + \epsilon_i$. For simplicity, all subject timecourses, $s$, have zero mean, we define $a$ as the norm of $g$ ($g^T g = a$), and we define

$\mathbb{E}[\epsilon_i^T \epsilon_i] = b$. If we have N subjects in total, the split half ISC ($shISC$) is:

$$\begin{aligned}
shISC &= \mathbb{E}\left[\rho\left(\sum_{i=1}^{N/2} s_i, \sum_{j=N/2+1}^{N} s_j\right)\right] \\
&= \mathbb{E}\left[\rho\left(\frac{N}{2}g + \sum_{i=1}^{N/2}\epsilon_i, \frac{N}{2}g + \sum_{j=N/2+1}^{N}\epsilon_j\right)\right] \\
&= \frac{\frac{N^2}{4}g^T g}{\sqrt{\frac{N^2}{4}g^T g + \mathbb{E}[\sum_{i=1}^{N/2}\epsilon_i^T\epsilon_i]}\sqrt{\frac{N^2}{4}g^T g + \mathbb{E}[\sum_{j=N/2+1}^{N}\epsilon_j^T\epsilon_j]}} \\
&= \frac{N^2 a}{N^2 a + 2Nb} \\
&= \frac{Nf}{Nf + 2} \qquad \text{(Equation 1)}
\end{aligned}$$

where $f = a/b$, or the signal to noise ratio (the ratio of norms between $g$ and $\epsilon_i$).

If we rearrange the variables in Equation 1, we find that:

$$f = \frac{2 * shISC}{-N * shISC - N}. \qquad \text{(Equation 2)}$$

We can thus calculate $pwISC$ and $looISC$ from $shISC$ as they relate to $f$ as follows. For $pwISC$ between subjects $i$ and $j$:

$$\begin{aligned}
pwISC &= \mathbb{E}[\rho(s_i, s_j)] \\
&= \mathbb{E}[\rho(g + \epsilon_i, g + \epsilon_j)] \\
&= \mathbb{E}\left[\frac{(g+\epsilon_i)^T(g+\epsilon_j)}{\sqrt{(g+\epsilon_i)^T(g+\epsilon_i)}\sqrt{(g+\epsilon_j)^T(g+\epsilon_j)}}\right] \\
&= \frac{g^T g}{\sqrt{g^T g + \mathbb{E}[\epsilon_i^T\epsilon_i]}\sqrt{g^T g + \mathbb{E}[\epsilon_j^T\epsilon_j]}} \\
&= \frac{a}{a+b} \\
&= \frac{f}{f+1},
\end{aligned}$$

and for $looISC$, between one left-out subject, and all others:

$$\begin{aligned}
looISC &= \mathbb{E}\left[\rho\left(s_1, \sum_{j=2}^{N} s_j\right)\right] \\
&= \mathbb{E}\left[\rho\left(g + \epsilon_1, (N-1)g + \sum_{j=2}^{N} \epsilon_j\right)\right] \\
&= \frac{(N-1)a}{\sqrt{a+b}\sqrt{(N-1)^2 a + (N-1)b}} \\
&= \frac{(N-1)f}{\sqrt{f+1}\sqrt{(N-1)^2 f + (N-1)}} \\
&= \frac{\sqrt{N-1}f}{\sqrt{f+1}\sqrt{(N-1)f+1}}.
\end{aligned}$$

Thus, for example, if there are 40 subjects, and $shISC$ is 0.5, the estimated $pwISC = 0.05$ and the estimated $looISC = 0.18$.
